# A prime editing strategy to rewrite the γ-globin promoters and reactivate fetal hemoglobin for sickle cell disease

**DOI:** 10.1101/2025.01.13.632780

**Authors:** Anne Chalumeau, Maria Bou Dames, Panagiotis Antoniou, Priyanka Loganathan, Margaux Mombled, Guillaume Corre, Martin Peterka, Mario Amendola, Carine Giovannangeli, Marcello Maresca, Annarita Miccio, Mégane Brusson

## Abstract

Fetal hemoglobin (HbF) reactivation is a promising therapy for β-hemoglobinopathies. We developed a prime editing strategy that introduces multiple mutations in the fetal γ-globin promoters of patients’ hematopoietic stem/progenitor cells (HSPCs), boosting HbF expression and offering alternative therapeutic perspectives.

---

Beta-hemoglobinopathies (sickle cell disease (SCD) and β-thalassemias) are anemias caused by mutations affecting the production of adult hemoglobin (Hb) in erythroid cells. Transplantation of autologous HSPCs genetically modified using CRISPR/Cas9 nuclease was approved as a treatment for β-hemoglobinopathies^1^; however, alternative strategies are desirable to improve safety and efficacy. Mutations in the fetal γ-globin (*HBG1/2*) promoters lead to hereditary persistence of HbF (HPFH) after birth, alleviating disease severity^2^. HPFH mutations either generate activator (KLF1^3^, TAL1^4^, GATA1^5^) binding sites (BSs) or disrupt repressor (BCL11A^6^, LRF^6^) BSs (**Fig.1A**). Disrupting repressor BSs via DNA double-strand break (DSB)-induced InDels using CRISPR/Cas9 nuclease reactivates HbF and corrects β-hemoglobinopathies^7^.

**Figure 1.**
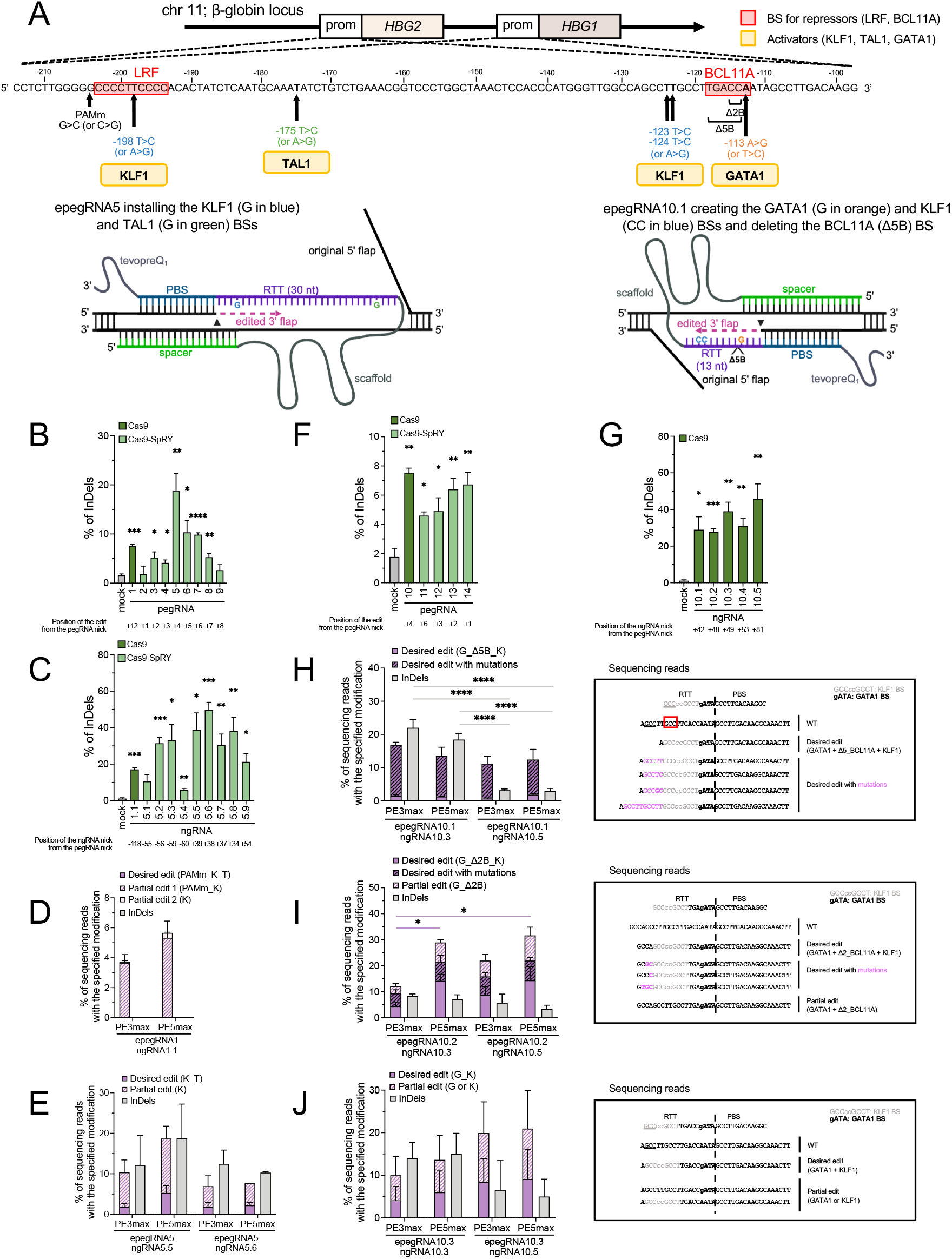
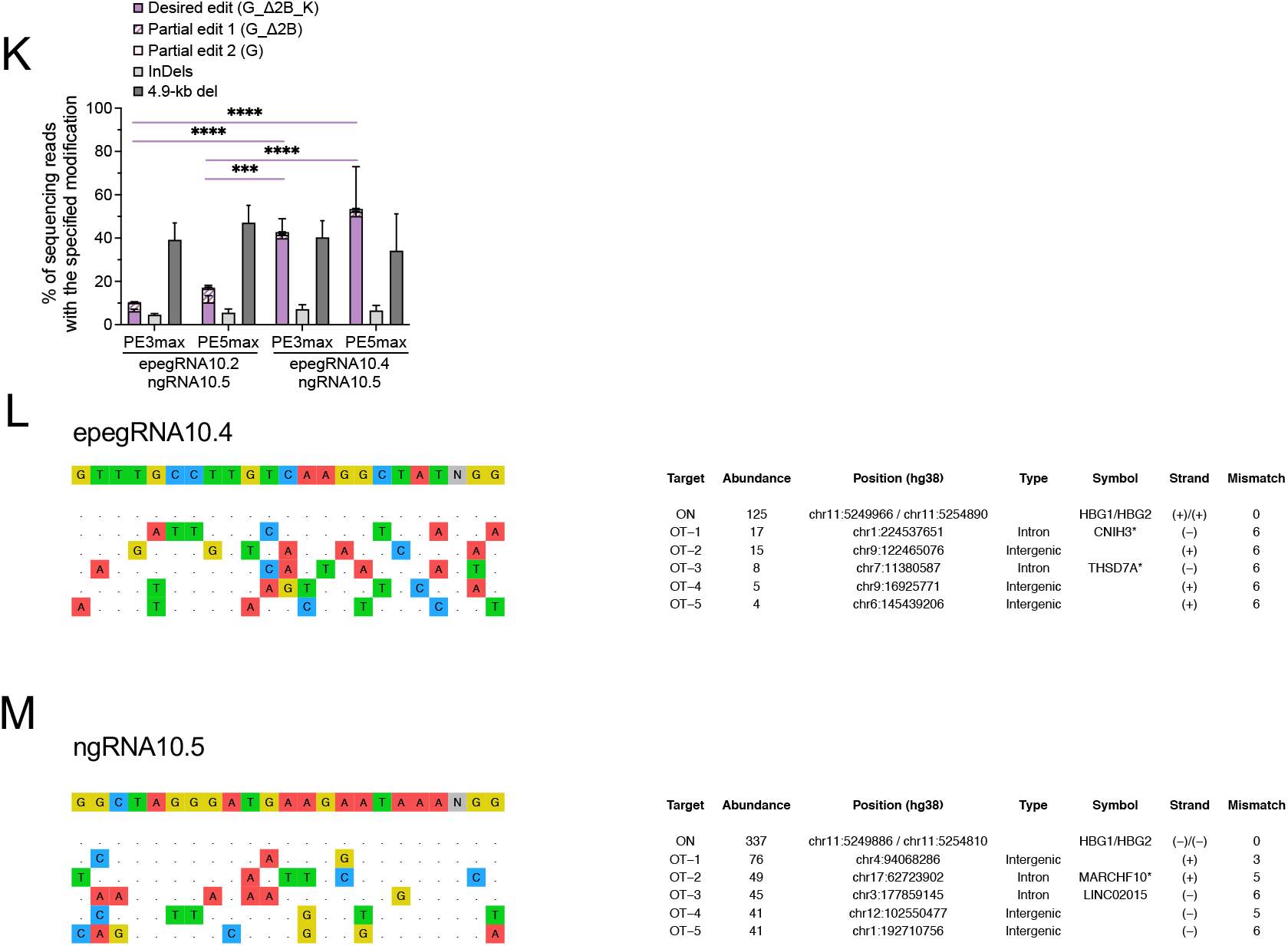
Screening of epegRNAs and ngRNAs targeting the *HBG1/2* promoters in K562 cells. **(A)** (top) Schematic representation of the β-globin locus on chromosome (chr) 11 including the *HBG1* and *HBG2* genes and their promoters (prom). Hereditary persistence of fetal hemoglobin (HPFH) or HPFH-like mutations (bold) disrupt LRF or BCL11A repressor binding sites (BSs; red boxes) or generate KLF1, TAL1, or GATA1 activator (yellow boxes) BSs. PAM-disrupting (PAMm) mutation and BCL11A BS-deletion (Δ2B: ΔCC, Δ5B: ΔTGACC) present in pegRNA1/epegRNA1, and in pegRNA10/epegRNA10.1/epegRNA10.2 are displayed bellow the promoter sequence. (bottom) Schematic representation of (left) epegRNA5 and (right) epegRNA10.1 targeting the *HBG1/2* promoter region. The epegRNA is composed (5’ to 3’) by a spacer, a scaffold, a primer binding site (PBS), a reverse transcription template (RTT) containing the edits (i.e., base substitutions and deletion indicated in colors and black), and the tevopreQ_1_ motif. Upon annealing of the spacer to the target DNA, the Cas9n induces a DNA single-strand break (i.e., nick, arrowhead) liberating the opposite strand (3’ flap), which anneals to the PBS of the epegRNA and primes the reverse transcription of the RTT. The interconversion of the original 5’ flap by the newly synthetized, edited 3’ flap and the degradation of the 5’ flap are essential steps to install the desired edits into the target region. **(B, C)** Frequency (%) of InDels generated by transfecting plasmids coding for (B) pegRNAs or (C) ngRNAs and Cas9 (pegRNA1 and ngRNA1.1) or Cas9-SpRY (pegRNA2 to pegRNA9 and ngRNA5.1 to ngRNA5.9) nucleases. While pegRNA1 was designed to install both KLF1, TAL1, and PAM-disrupting (PAMm) mutations (to avoid retargeting of the region), pegRNA2 to pegRNA5 have a shorter RTT installing only the KLF1 BS **(D)** Percentage of NGS reads containing the desired edit (3 mutations: PAM-disrupting mutation, KLF1 and TAL1 BSs; PAMm_K_T), the partial edits (1 or 2 mutations; K or PAMm_K) or InDels generated using epegRNA1, ngRNA1.1 and PE3max or PE5max in the top 30% GFP+ cells. **(E)** Percentage of NGS reads containing the desired edit (2 mutations: KLF1 and TAL1; K_T), the partial edit (1 mutation; K) or InDels generated using epegRNA5, with ngRNA5.5 (left) or ngRNA5.6 (right) and PE3max or PE5max (in their PAM-less SpRY version) in the top 30% GFP+ cells. **(F, G)** Frequency (%) of InDels generated by transfecting plasmids coding for (F) pegRNAs or (G) ngRNAs and Cas9 (pegRNA10 and ngRNA10.1 to ngRNA10.5) or Cas9-SpRY (pegRNA11 to pegRNA14) nucleases. (left) Percentage of NGS reads containing the desired edit (3 modifications: GATA1 BS, BCL11A-5-nt BS deletion [ΔTGACC from -114 to -118 upstream of *HBG1/2* TSS], KLF1 BS: G_Δ5B_K), the desired edit with mutations, or InDels induced by epegRNA10.1 with ngRNA10.3 or ngRNA10.5 in the top 30% GFP+ cells. (right) The sequences of the PBS and RTT of epegRNA10.1 are aligned on the most frequent NGS reads. The KLF1 and GATA1 BSs are indicated in grey and bold, respectively, and the desired base conversions in lower case. The 5bp-deletion of the edited 3’-flap facilitates the annealing of its GCC trinucleotide (underlined) to the closest complementary motif (red square) leading to desired edit with mutations (pink) +/-scaffold incorporation (pink and bold). **(I)** (left) Percentage of NGS reads containing the desired edit (3 modifications: GATA1 BS, BCL11A-BS 2-nt deletion [ΔCC -117/-118 upstream of *HBG1/2* TSS], KLF1 BS: G_Δ2B_K), the desired edit with mutations, the partial edit (G_Δ2B) or InDels induced by epegRNA10.2 with ngRNA10.3 or ngRNA10.5 in the top 30% GFP+ cells. (right) The sequences of the PBS and RTT of epegRNA10.2 are aligned on the most frequent NGS reads. The KLF1 and GATA1 BSs are indicated in grey and bold, respectively, and the desired base conversions in lower case. **(J)** (left) Percentage of NGS reads containing the desired edit (2 modifications: GATA1 and KLF1 BSs: G_K), the partial edits (GATA1 or KLF1 BSs), or InDels induced by epegRNA10.3 with ngRNA10.3 or ngRNA10.5 in the top 30% GFP+ cells. (right) The sequences of the PBS and RTT of epegRNA10.3 are aligned on the most frequent NGS reads. The KLF1 and GATA1 BSs are indicated in grey and bold, respectively, and the desired base conversions in lower case. The long 3’-flap facilitates the annealing of its GCC trinucleotide (underlined) to the expected complementary motif preventing the formation of desired edit with unintended mutations. **(K)** Percentage of NGS reads containing the desired edits, the partial edits or InDels and frequency (%) of 4.9-kb deletion measured by ddPCR (as previously described^16^ using the primers and probes described in **Table S2**) generated using epegRNA10.2 or epegRNA10.4 with ngRNA10.5 into the *HBG1/2* promoters in the top 30% GFP+ cells. (**L, M**) Top five (M) epegRNA10.4- and (N) ngRNA10.5-dependent off-target DNA sites, as identified by GUIDE-seq^16^ in K562 cells. epegRNA and ngRNA were coupled with a Cas9 nuclease equivalent to the Cas9 nickase included in the prime editor except for carrying the wild-type amino acid allowing nuclease activity. The protospacer targeted by each epegRNA or ngRNA and the PAM are reported in the first line, followed by the on- and off-target sites and their mismatches with the on-target (highlighted in color). For each target, the abundance (i.e., the number of cells containing the target site), the chromosomal coordinates (Human GRCh38/hg38), the type of region, the gene and the targeted strand, and the number of mismatches with the epegRNA or ngRNA protospacer are reported. In total, we identified 14 off-targets for epegRNA10.4, with only 5 with an abundance >3, and 44 off-targets for ngRNA10.5, with 27 with an abundance >3. Most of them have a high number of mismatches (≥5) and map to non-exonic regions. The datasets supporting the GUIDE-seq results of this article are available for the reviewers in the BioProject repository under the BioProject ID: PRJNA1200278 (http://www.ncbi.nlm.nih.gov/bioproject/1200278). (**B, C, F, G**) Samples mock-transfected with TE (Tris-EDTA) buffer were used as controls. PCR products were subjected to Sanger sequencing (as previously described^16^ sing the primers and probes described in **Table S2**) and InDels were measured using the TIDE software^26^. Bars represent the mean ± SEM of 3 biologically independent replicates. Statistical significance was assessed between mock and treated samples using an unpaired t-test and displayed for ****p<0.0001, ***p<0.001, **p<0.01, *p<0.05. (**D, E, H-L**) NGS analysis were performed as previously described^16^ using the primers and probes described in **Table S2**. A customized Python pipeline was used to align NGS reads to a reference amplicon sequence and count desired and partial edits, and other InDels. The “Desired edit” and “Partial edits” categories do not contain InDels and are stacked. InDels are located at the nick induced by the epegRNA or the ngRNA. Bars represent the mean ± SEM of n= (D, E, H-J) 3 or (K, L) 3-4 biologically independent replicates. Statistical significance was assessed using two-way ANOVA with multiple comparisons: ****p<0.0001, ***p<0.001, *p<0.05, or not significative.

Base and prime editors allow the insertion of precise edits. While base editors are limited to base transitions, prime editors (PEs) enable all base conversions, insertions, deletions, and combinations of these mutations^8^. The latter is a desirable approach for β-hemoglobinopathies as co-occurrence of multiple HPFH mutations in a healthy individual led to higher HbF than individual mutations^9^. PEs contain a *Sp*Cas9 nickase (Cas9n) fused to a reverse transcriptase (RT). The prime editing guide RNA (pegRNA) harbours a scaffold binding Cas9n, a spacer driving PE to the target (protospacer) where Cas9n induces a single-strand break (“nick”) upstream of the protospacer adjacent motif (PAM), a primer binding site (PBS) annealing to the nicked target region and serving as a template for the RT and an RT-template (RTT) containing the desired edits (DEs; **Fig.1A**). The RTT is reverse-transcribed forming the edited 3’ flap and the non-edited 5’ flap is removed (PE2 system). To increase the editing efficiency, PE2 was improved by using a nicking guide RNA (ngRNA) that nicks the non-edited strand (that is then repaired using the edited strand as template; PE3^8^) and a dominant negative MLH1 (MLH1dn) that inhibits the mismatch DNA repair pathway (MMR; PE5), which reincorporates the original nucleotides in the edited strand^10^.

Here, we used prime editing to introduce in *HBG1/2* promoters multiple HPFH/HPFH-like mutations^5,6,11^ in cell lines and primary patients’ HSPCs.

We first designed pegRNAs to generate KLF1 (-198 T>C) and TAL1 (-175 T>C) BSs into the *HBG1/2* promoters (**Fig.1A, TableS1**). These pegRNAs are compatible with PEmax or PEmax-SpRY containing the NGG-PAM Cas9n or the near PAM-less Cas9n-SpRY, respectively^10^. As prime-editing efficiency correlates with Cas9-induced InDel frequency^12^, we screened pegRNAs with Cas9 in K562 cells. We selected pegRNA1 and pegRNA5 showing the highest InDel frequency, and ngRNA1.1, ngRNA5.5 and ngRNA5.6 (**Fig.1B-C, TableS1**). Then, we generated epegRNA1 and epegRNA5, which contains a 36-nt and 30-nt RTT inserting KLF1/TAL1 BSs, respectively (**Fig.1A, TableS1**), and tevopreQ_1_ that enhances pegRNA stability^13^. epegRNA1 also installs a PAM-disrupting (PAMm) mutation (to avoid retargeting of the region; **Fig.1A**). These epegRNAs were electroporated with PE3/5max- and GFP-expressing plasmids in K562 cells and editing was assessed. While epegRNA1 installed only the KLF1 BS and PAMm, epegRNA5 generated simultaneously KLF1/TAL1 BSs, although at a low frequency (even when using PE5) and with significant InDel generation (that can potentially occur using PEs^14,15^; **Fig.1D-E**). These results suggest that the low RT processivity limits the installation of DEs located far from the nick (i.e., the TAL1 BS) using pegRNA with long RTT (i.e., ≥30 nt). This prompted us to design pegRNAs with shorter RTT (i.e., 11 to 14 nt) to insert HPFH mutations that are both close to the nick in the -115 of the *HBG1/2* promoters. In particular, we simultaneously inserted GATA1 (-113 A>G) and KLF1 (-123/-124 T>C) BSs and deleted the BCL11A BS (5-nt deletion that also disrupts the PAM: Δ5B, ΔTGACC) (**Fig.1A, TableS1**). We selected the pegRNA10 (RTT of 13 nt), ngRNA10.3 and ngRNA10.5 showing the highest InDel frequency, and added the tevopreQ_1_ to generate epegRNA10.1 (**Fig.1A,F-G, TableS1**). In K562 cells, we observed a low frequency of DEs, and the generation of DEs with additional mutations (**Fig.1H**). This could be due to the Δ5B that reduces the complementarity of the edited 3’-flap to the opposite strand, facilitating the annealing of the GCC trinucleotide of the 3’-flap to the closest complementary motif. Moreover, some of these unintended mutations contain a partial incorporation of the scaffold that is reverse-transcribed by the RT. ngRNA10.3 caused significantly higher InDel frequencies than ngRNA10.5 (**Fig.1H**). To increase 3’-flap complementarity, we designed epegRNA10.2 and epegRNA10.3 containing the epegRNA10.1 spacer and PBS, and a 16-or 18-nucleotide-long RTT inserting the GATA1/KLF1 BSs and deleting or not the BCL11A BS 2-nt core (Δ2B:ΔCC; **Fig.1A, TableS1**)^6^. This modification strongly favored DEs over unintended flap annealing (**Fig.1I-J**). Scaffold incorporation, which should not impact the promoter activity, still occurred with the epegRNA10.2, but always in combination with DEs (**Fig.1I**). The epegRNA10.2/ngRNA10.5 combination was the most efficient, reaching 34% of total prime-editing events (when using PE5), with the lowest InDel frequency (**Fig.1J**). Of note, epegRNA10.2 and epegRNA10.3 generated partial edits suggesting low processivity of the PEmax leading to incomplete reverse transcription of the ≥16-nucleotide-long RTT or exonuclease-mediated 3’-flap resection (**Fig.1I-J**).

Therefore, we elongated the epegRNA10.2 RTT (+7nt, epegRNA10.4) to prevent 3’-flap resection, and further favor the correct annealing of the edited 3’ flap with the aim to increase DE installation (**Fig.1K**). In K652 cells, epegRNA10.4 led to a 6.4-fold higher editing efficiency than epegRNA10.2 (**Fig.1K**). We also observed the *HBG1*-*HBG2* 4.9-kb deletion, likely due to DSB generation^14^ or strand displacement because of the generation of two nicks in *cis*^14,16,17^(**Fig1.K-L**). Noteworthy, for the most promising epegRNA10.4/ngRNA10.5 combination, GUIDE-seq analysis^16^ revealed few off-targets with ≥ 3 mismatches, which mapped to non-exonic regions, suggesting that if any, off-target activity will not dramatically impact gene/protein expression (**Fig1.M-N**).

Since MMR genes are poorly expressed in HSPCs^10,18^ (**Fig.2A**), we tested epegRNA10.4/ngRNA10.5 in SCD HSPCs using PE3max or Cas9 nuclease-PEmax (PEnmax) that can increase prime-editing efficiency^19^ (**Fig.2B**). Both PEs inserted DEs in HSPCs with variable efficiencies, as previously reported^18^. However, PEnmax outperformed PE3max reaching 7% of desired/partial edits (**Fig.2C**). Both systems also generated InDels and 4.9-kb deletions, with a higher frequency for the Cas9 nuclease-based PEnmax (**Fig.2C-D**).

**Figure 2.**
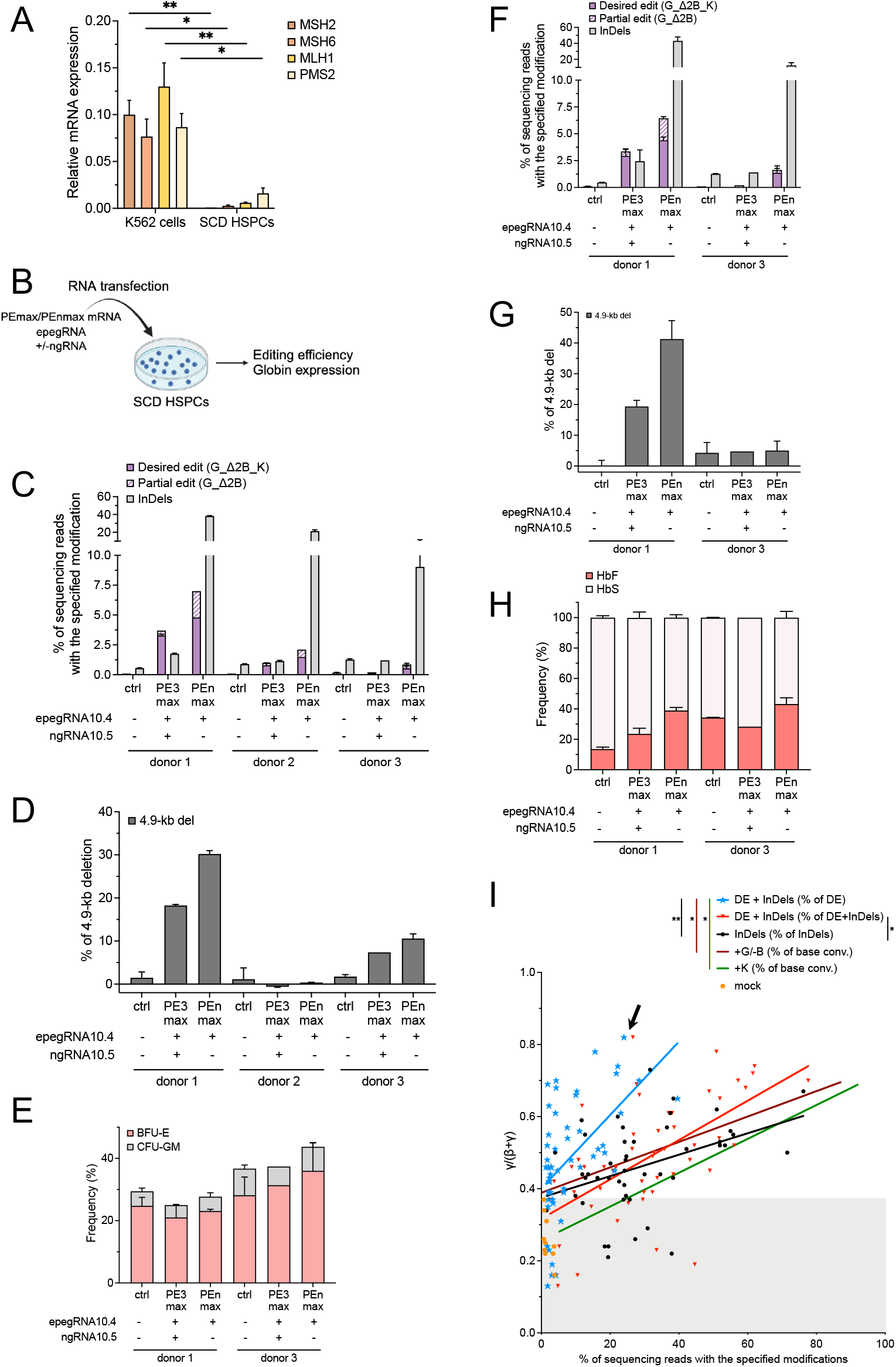
BCL11A BS disruption combined with generation of GATA1 and KLF1 BSs in the *HBG1/2* promoters reactivates HbF in erythroid cells differentiated from SCD HSPCs. **(A)** Relative mRNA expression of genes encoding the main factors involved in MMR in K562 cells and peripheral blood CD34+ HSPCs from 3 SCD donors, as measured by RT-qPCR as previously described^10^. *MSH2, MSH6, MLH1*, and *PSM2* mRNA expression was normalized to the expression of *ACTB* mRNA. Bars represent the mean ±SEM of n=3 biologically independent replicates for K562 cells and n=3 donors for SCD HSPCs. Statistical significance was assessed using an unpaired t-test: **p<0.01, and *p<0.05. **(B)** Experimental protocol used for prime editing experiments in peripheral blood CD34+ HSPCs from SCD donors. The PEmax or the PEnmax mRNA (6.4 μg of mRNA *in vitro* transcribed as previously described^16^ using the AM1334 or AM1345 kit, Thermo Fisher), epegRNA (200 pmol, IDT), +/-the ngRNA (200 pmol, Synthego), and the GFP mRNA (0.5 μg, TebuBio) were co-transfected [P3 Primary Cell 4D-Nucleofector X Kit (Lonza) and the CA137 program (Nucleofector 4D)] in SCD HSPCs 24 h after cell thawing and culture in pre-activation medium [PAM: StemSpan (STEMCELL Technologies) supplemented with penicillin/streptomycin (ThermoFisher) and L-glutamine (ThermoFisher), 750 μM StemRegenin1 (STEMCELL Technologies), and the following recombinant human cytokines (PeproTech): human stem cell factor (SCF; 300 ng/mL), Flt-3L (300 ng/mL), thrombopoietin (TPO; 100 ng/mL), and interleukin-3 (IL-3; 60 ng/mL)]. After transfection, HSPCs were either cultured in PAM or plated in a semi-solid medium to induce erythroid differentiation (colony forming assay). Prime editing efficiency was evaluated in HSPCs 6 days post-transfection and in BFU-Es by NGS (Illumina, as previously described^16^ using the primers and probes described in **Table S2**) and a customized Python pipeline that aligns NGS reads to a reference amplicon sequence and counts precise and partial edits and InDels (at the pegRNA or ngRNA nick sites). Globin mRNA expression was quantified in single BFU-E by RT-qPCR and Hb expression in BFU-E pools (25 colonies) by HPLC, as previously described^27^. **(C)** Percentage of NGS reads containing the desired edits (3 modifications: GATA1 BS, BCL11A-BS 2bp deletion [ΔCC -117/-118 upstream of *HBG1/2* TSS], KLF1 BS: G_Δ2B_K), the partial edit (G_Δ2B) or InDels in HSPCs from 3 SCD donors using epegRNA10.4 +/-ngRNA10.5 and PE3max or PEnmax **(D)** Frequency (%) of 4.9-kb deletion measured by ddPCR generated using epegRNA10.4 +/-ngRNA10.5 in HSPCs from 3 SCD donors. **(E)** Frequency (%) of BFU-E and CFU-GM derived from ctrl- and PE-transfected HSPCs. Results are represented as frequency of colonies obtained from 500 plated HSPCs and shown as mean±SEM. ns, two-way ANOVA test. **(F)** Percentage of NGS reads containing the desired edits (3 modifications: GATA1 BS, BCL11A-BS 2bp deletion [CC -117/-118 upstream of *HBG1/2* TSS], KLF1 BS: G_Δ2B_K), the partial edit (G_Δ2B) or InDels in SCD bulk BFU-Es using epegRNA10.4 +/-ngRNA10.5 and PE3max or PEnmax. **(G)** Frequency (%) of 4.9-kb deletion measured by ddPCR in SCD bulk BFU-Es using epegRNA10.4 +/-ngRNA10.5 and PE3max or PEnmax. **(H)** HbF and sickle Hb (HbS) expression measured by HPLC^27^ in SCD BFU-E pools treated with epegRNA10.4 +/-ngRNA10.5 and PE3max or PEnmax. **(I)** γ-globin mRNA expression (RT-qPCR, normalized on α-globin mRNA as previously described^27^) in individual BFU-Es derived from SCD HSPCs (1 donor). HSPCs were mock-transfected or transfected with PEn and epegRNA10.4. Each single BFU-E was classified depending on its genotype: “DE + InDels” (n=53 colonies containing the desired edit G_Δ2B_K and InDels) for which we reported only the % of DE (blue) or the % of total edit (DE + InDels; red), “InDels” (n= 46 colonies containing only InDels; black), and “+G/-B” or “+K” (n=15 and n= 16 colonies, containing the GATA1 BS and a disrupted BCL11A BS or the KLF1 BS, generated using the ABE8-13m^28^ and published gRNAs^11,29^ for which we reported the frequency of base conversion (conv.). Statistical significance between two curves was assessed using linear regression: **p<0.01, *p<0.05, or not significant. The arrow indicates a single colony containing 24% of DE with minimal InDels and expressing the highest γ-globin level. (**C-H**) Bars represent the mean ± SEM of n=1-3 technical replicates per donor (3 donors for C, D and 2 donors for E-H). Tris-EDTA buffer-transfected samples and cells transfected with PEmax or PEnmax and no pegRNA/ngRNA were used as controls (ctrl). (**C, F**) The “Desired edits” and “Partial edits” categories do not contain InDels and are stacked. InDels are located at the nick induced by the epegRNA or the ngRNA.

We then evaluated if multiple HPFH/HPFH-like mutations induce high HbF. Twenty-four hours post-transfection, HSPCs were plated in a semi-solid medium promoting the growth of erythroid clones (BFU-E). PEs did not affect growth and multilineage differentiation of hematopoietic progenitors (**Fig.2E**). Editing efficiency (including 4.9-kb deletion frequency) were similar between BFU-E and HSPCs, demonstrating no counterselection of edited erythroid cells (**Fig.2C-G**). PE3max/PEnmax treatment led to HbF reactivation compared to controls (**Fig.2H**). Importantly, a clonal analysis showed that that the BFU-Es carrying DE and InDels express significantly higher γ-globin levels than colonies carrying only InDels or individual mutations (**Fig.2I**).

In conclusion, we provided proof-of-concept for combining multiple HPFH/HPFH-like mutations to boost HbF expression. This could give a better selective advantage to erythroid cells, thus requiring a lower fraction of edited HSPCs to achieve a therapeutic benefit^20^ (a scenario that is possible with an *in vivo* HSPC-targeting strategy^21^). However, we highlighted the limited efficiency of the current prime-editing systems for long pegRNAs and hard-to-edit loci^22^. The use of PEnmax increased the incorporating of DEs, but unintended events, such as InDels, should be minimized e.g., by using DNA-PK inhibitors^23^. Further optimization could be performed to increase prime editing efficiency^24,25^. Overall, these results advance the understanding of the repair mechanisms involved in prime-editing strategies and could lead to an effective, safe and universal treatment of β-hemoglobinopathies (independently from the disease-causing mutation).

## Acknowledgements

We thank Dr. Sandra Manceau for the collection of the blood samples and the flow cytometry platform of the SFR Necker. This work was supported by state funding from the French National Research Agency (Agence Nationale de la Recherche) (ANR-10-IAHU-01 and ANR-22-CE17-0028 PEMGeT), the European Commission (HORIZON-PathFinder EdiGenT grant no. 101070903), the AFM-Telethon (PhD fellowship grant no. 23879, Research Grants no. 24331, and 25179), and the COST (European Cooperation in Science and Technology (the COST Action Gene Editing for the treatment of Human Diseases, CA21113). This study was also supported by the EUR G.E.N.E. (reference #ANR-17-EURE-0013) and is part of the Université Paris Cité IdEx #ANR-18-IDEX-0001 funded by the French Government through its “Investments for the Future” program.

## Author contributions

A.C. designed, conducted experiments, analyzed the data, and wrote the paper. M.B.D. designed, conducted experiments and analyzed the data. P.L. conducted experiments and analyzed data. M.M., G.C. and M.A. designed, conducted and analyzed the GUIDE-seq experiment. C.G. analyzed NGS data and contributed to the design of the experimental strategy. P.A., M.P., and M.M conducted NGS experiments and contributed to the design of the experimental strategy. A.M. and M.B. conceived the study, designed experiments, and wrote the paper.

