## Supplemental Table S1, S2 for "A prime editing strategy to rewrite the γ-globin promoters and reactivate fetal hemoglobin for sickle cell disease"

**Table S1: List of pegRNAs and ngRNAs sequences targeting the *HBG1/2* promoters.**

| Name | spacer 5'-3' | scaffold 5'-3' | RTT 5'-3' | PBS 5'-3' | linker 5'-3' | tevopreQ1 motif 5'-3' | total length |
|---|---|---|---|---|---|---|---|
| pegRNA1 | AAGCAGCAGTATCCTCTTGG | GTTTTAGAGCTAGAAATAGCAAGTTAAAATAAGGCTAGTCCGTTATCAACTTGAAAAAGTGGCACCGAGTCGGTGC | TGTTTGCATTGAGATAGTGTGGGGGAGGGGGCCCCA | AGAGGATACTGCT | | | 145 |
| pegRNA2 | TCCTCTTGGGGGCCCCTTCC | GTTTTAGAGCTAGAAATAGCAAGTTAAAATAAGGCTAGTCCGTTATCAACTTGAAAAAGTGGCACCGAGTCGGTGC | AGATAGTGTGGGGG | AGGGGCCCCCAAG | | | 123 |
| pegRNA3 | ATCCTCTTGGGGGCCCCTTC | GTTTTAGAGCTAGAAATAGCAAGTTAAAATAAGGCTAGTCCGTTATCAACTTGAAAAAGTGGCACCGAGTCGGTGC | GATAGTGTGGGGGA | GGGGCCCCCAAGA | | | 123 |
| pegRNA4 | TATCCTCTTGGGGGCCCCTT | GTTTTAGAGCTAGAAATAGCAAGTTAAAATAAGGCTAGTCCGTTATCAACTTGAAAAAGTGGCACCGAGTCGGTGC | ATAGTGTGGGGGAG | GGGCCCCCAAGAG | | | 123 |
| pegRNA5 | GTATCCTCTTGGGGGCCCCT | GTTTTAGAGCTAGAAATAGCAAGTTAAAATAAGGCTAGTCCGTTATCAACTTGAAAAAGTGGCACCGAGTCGGTGC | TAGTGTGGGGGAGG | GGCCCCCAAGAGG | | | 123 |
| pegRNA6 | AGTATCCTCTTGGGGGCCCC | GTTTTAGAGCTAGAAATAGCAAGTTAAAATAAGGCTAGTCCGTTATCAACTTGAAAAAGTGGCACCGAGTCGGTGC | AGTGTGGGGGAGGG | GCCCCCAAGAGGA | | | 123 |
| pegRNA7 | CAGTATCCTCTTGGGGGCCC | GTTTTAGAGCTAGAAATAGCAAGTTAAAATAAGGCTAGTCCGTTATCAACTTGAAAAAGTGGCACCGAGTCGGTGC | GTGTGGGGGAGGGG | CCCCCAAGAGGAT | | | 123 |
| pegRNA8 | GCAGTATCCTCTTGGGGGCC | GTTTTAGAGCTAGAAATAGCAAGTTAAAATAAGGCTAGTCCGTTATCAACTTGAAAAAGTGGCACCGAGTCGGTGC | TGTGGGGGAGGGGC | CCCCAAGAGGATA | | | 123 |
| pegRNA9 | AGCAGTATCCTCTTGGGGGC | GTTTTAGAGCTAGAAATAGCAAGTTAAAATAAGGCTAGTCCGTTATCAACTTGAAAAAGTGGCACCGAGTCGGTGC | GTGGGGGAGGGGCC | CCCAAGAGGATAC | | | 123 |
| epegRNA1 | AAGCAGCAGTATCCTCTTGG | GTTTCAGAGCTATGCTGGAAACAGCATAGCAAGTTGAAATAAGGCTAGTCCGTTATCAACTTGAAAAAGTGGCACCGAGTCGGTGC | TGTTTGCATTGAGATAGTGTGGGGGAGGGGGCCCCA | AGAGGATACTGCT | ATAAATAT | CGCGGTTCTATCTAGTTACGCGTTAAACCAACTAGAA | 200 |
| epegRNA5 | GTATCCTCTTGGGGGCCCCT | GTTTCAGAGCTATGCTGGAAACAGCATAGCAAGTTGAAATAAGGCTAGTCCGTTATCAACTTGAAAAAGTGGCACCGAGTCGGTGC | GATGTTTGCATTGAGATAGTGTGGGGGAGG | GGCCCCCAAGAGG | CGAGAAGT | CGCGGTTCTATCTAGTTACGCGTTAAACCAACTAGAA | 194 |

| Name | spacer 5'-3' | scaffold | RTT 5'-3' | PBS 5'-3' | linker 5'-3' | tevopreQ1 motif 5'-3' | total length |
|---|---|---|---|---|---|---|---|
| pegRNA10 | GTTTGCCTTGTCAAGGCTAT | GTTTTAGAGCTAGAAATAGCAAGTTAAAATAAGGCTAGTCCGTTATCAACTTGAAAAAGTGGCACCGAGTCGGTGC | GCCCCGCCTGATA | GCCTTGACAAGGC | | | 122 |
| pegRNA11 | TGCCTTGTCAAGGCTATTGG | GTTTTAGAGCTAGAAATAGCAAGTTAAAATAAGGCTAGTCCGTTATCAACTTGAAAAAGTGGCACCGAGTCGGTGC | GCCAGCCCCGCCTG | ATAGCCTTGACAA | | | 123 |
| pegRNA12 | TTGCCTTGTCAAGGCTATTG | GTTTTAGAGCTAGAAATAGCAAGTTAAAATAAGGCTAGTCCGTTATCAACTTGAAAAAGTGGCACCGAGTCGGTGC | AGCCCCGCCTGA | TAGCCTTGACAAG | | | 121 |
| pegRNA13 | TTTGCCTTGTCAAGGCTATT | GTTTTAGAGCTAGAAATAGCAAGTTAAAATAAGGCTAGTCCGTTATCAACTTGAAAAAGTGGCACCGAGTCGGTGC | GCCCCGCCTGAT | AGCCTTGACAAGG | | | 121 |
| pegRNA14 | AGTTTGCCTTGTCAAGGCTA | GTTTTAGAGCTAGAAATAGCAAGTTAAAATAAGGCTAGTCCGTTATCAACTTGAAAAAGTGGCACCGAGTCGGTGC | TTGCCTGATAG | CCTTGACAAGGCA | | | 120 |
| epegRNA10.1 | GTTTGCCTTGTCAAGGCTAT | GTTTCAGAGCTATGCTGGAAACAGCATAGCAAGTTGAAATAAGGCTAGTCCGTTATCAACTTGAAAAAGTGGCACCGAGTCGGTGC | GCCCCGCCTGATA | GCCTTGACAAGGC | TCCTAATC | CGCGGTTCTATCTAGTTACGCGTTAAACCAACTAGAA | 177 |
| epegRNA10.2 | GTTTGCCTTGTCAAGGCTAT | GTTTCAGAGCTATGCTGGAAACAGCATAGCAAGTTGAAATAAGGCTAGTCCGTTATCAACTTGAAAAAGTGGCACCGAGTCGGTGC | GCCCCGCCTTGAGATA | GCCTTGACAAGGC | TCCTTATC | CGCGGTTCTATCTAGTTACGCGTTAAACCAACTAGAA | 182 |
| epegRNA10.3 | GTTTGCCTTGTCAAGGCTAT | GTTTCAGAGCTATGCTGGAAACAGCATAGCAAGTTGAAATAAGGCTAGTCCGTTATCAACTTGAAAAAGTGGCACCGAGTCGGTGC | GCCCCGCCTTGACCGATA | GCCTTGACAAGGC | CTCTTCTA | CGCGGTTCTATCTAGTTACGCGTTAAACCAACTAGAA | 180 |
| epegRNA10.4* (used in K562 cells) | GTTTGCCTTGTCAAGGCTAT | GTTTCAGAGCTATGCTGGAAACAGCATAGCAAGTTGAAATAAGGCTAGTCCGTTATCAACTTGAAAAAGTGGCACCGAGTCGGTGC | TTGGCCAGCCCCGCCTTGAGATA | GCCTTGACAAGGC | TCCGGTAA | CGCGGTTCTATCTAGTTACGCGTTAAACCAACTAGAA | 187 |
| epegRNA10.4** (used in HSPCs) | GTTTGCCTTGTCAAGGCTAT | GTTTTAGAGCTAGAAATAGCAAGTTAAAATAAGGCTAGTCCGTTATCAACTTGAAAAAGTGGCACCGAGTCGGTGC | TTGGCCAGCCCCGCCTTGAGATA | GCCTTGACAAGGC | TCCGGTAA | CGCGGTTCTATCTAGTTACGCGTTAAACCAACTAGAA | 187 |

** and **: the epegRNA differ by the scaffold sequence, * was used in K562 cells, and ** in HSPCs and GUIDE-seq analysis*

*The scaffold sequence (GTTTTAGAGCTAGAAATAGCAAGTTAAAATAAGGCTAGTCCGTTATCAACTTGAAAAAGTGGCACCGAGTCGGTGC) is from Dang et. Al, Genome Biol., 2015 and was selected to improve Cas9 efficiency*

*The scaffold sequence (GTTTCAGAGCTATGCTGGAAACAGCATAGCAAGTTGAAATAAGGCTAGTCCGTTATCAACTTGAAAAAGTGGCACCGAGTCGGTGC) is from Anzalone et. Al, Nature, 2019 for efficient prime editing*

**Table S2: primers/probes used for sequencing analysis and ddPCR**

| **Sanger sequencing** | | |
|---|---|---|
| **Amplified region** | **F/R** | **Sequence (5' to 3')** |
| *HBG1/2* promoter | F | AAAAACGGCTGACAAAAGAAGTCCTGGTAT |
| | R | ATAACCTCAGACGTTCCAGAAGCGAGTGTG |

F, forward primer; R, reverse primers

| **Deep sequencing for prime editing experiments** | | | |
|---|---|---|---|
| **Amplified region** | **target epegRNA** | **F/R** | **Sequence (5' to 3')** |
| HBG1/2 promoter | epegRNA1 | F | CGGCTGACAAAAGAAGTCCT |
| | | R | TTTAGCCAGGGACCGTTTCAG |
| | epegRNA5 | F | GGAATGACTGAATCGGAACAAGG |
| | | R | CTGGCCTCACTGGATACTCT |
| | epegRNA10.1/10.2/10.3/10.4 | F | AAACGGTCCCTGGCTAAACT |
| | | R | CCAGAAGCGAGTGTGTGGAA |

F, forward primer; R, reverse primers

| **ddPCR analysis** | | | |
|---|---|---|---|
| | F/R | Sequence (5' to 3') | Flurophore |
| primers | F | AAAGAGAGGTGGAAATGAGG | |
| | R | CCACTTTGACTGAGCCAATA | |
| probe | HBG1 | CCAGTAGAAAGAACTTTCATCTTTCCCTC | FAM |
| probe | HBG2 | CTTTCCCCTATTTTTGTTATTCGTTTTA | VIC |

F, forward primer; R, reverse primers
